# Different autotrophic enrichments yield efficient inocula for biocathodic applications

**DOI:** 10.1101/2021.01.21.427587

**Authors:** Arda Gülay, Marlene Mark Jensen, Thomas Sicheritz-Pontén, Barth F. Smets

**Affiliations:** Harvard University; Denmark Technical University; University of Copenhagen; Technical University of Denmark

## Abstract

The uptake of extracellular electrons from cathodes by microbes is a recently discovered phenomenon. However, current knowledge on the diversity of microbes accepting extracellular electrons and populating biocathodes is scarce. Research is required to explore the distribution of extracellular electron uptake metabolism in the tree of life across microbial guilds. Here we characterize the electron uptake ability of microbial guilds enriched on H_2_, S_2_O_3_^2-^, or CH_4_ and NH_4_^+^ from the same inoculum taken from a groundwater treatment sand-filter. We hypothesized that functional microbes having dense outer membrane cytochromes in their native pathway, such as pathways of NH_4_^+^, CH_4_, S_2_O_3_^2-^ and H_2_, can perform extracellular electron uptake. We aimed of addressing the following questions: (1) Are there any known microbial member of anticipated function performing extracellular electron transfer? (2) How does electron uptake efficiency vary between the different functional guilds? (3) How is the dissipated electron energy distributed across metabolisms? We developed and applied a novel pipeline to identify taxa utilizing direct electron energy and utilizing secondary microbial products. We report the putative direct electron uptake metabolism of types belonging to *Methylomonas, UBA6140*, and *Nitrosomonas*. Furthermore, members of *Streptococcaceae, Rhizobiaceae, Streptococcus, Brevundimonas, Chryseobacterium*, and *Pseudomonas* are detected as electroactive taxa. Our results reveal novel insights into the diversity, electrochemical activity, and metabolism of taxa performing direct electron uptake.

## Introduction

The uptake of extracellular electrons by microbes is a recently discovered microbial energetic pathway (Bond et al., 2002). Electron uptake process was successfully applied to bioremediate metals and organic contaminants (Zhang et al., 2013) or for microbial electrosynthesis to produce fuels and chemicals or to catalyze processes such as reduction of oxygen, CO_2_, nitrate (Rabaey and Rozendal, 2010; Huang et al., 2011; Marshall et al., 2012; Lovley and Nevin, 2013).

Microbes were first demonstrated to catalyze anodic organic carbon oxidation through donation of electrons to the anode (Kim et al., 1999; Rabaey et al., 2004; Holmes et al., 2006; Richter et al., 2009). Gregory et al. (2004) and later Rabaey et al. (2008) showed that pure cultures of bacteria could take up electrons and grow at a cathode. Extracellular electron transfer (EET) at cathodes has since been demonstrated for both single microbial strains, such as *acetogens* (Kracke et al., 2015), *methanogenic archaeon* (Beese-Vasbender et al., 2015), *Acidithiobacillus* (Carbajosa et al., 2010), *Acinetobacter* (Freguia et al., 2010), *Mariprofundus* (Summers et al., 2013) and *Rhodopseudomonas* (Doud and Angenent, 2014), and microbial mixed communities harvested from marine sediments (Bond et al., 2002) and activated sludge (Zhang et al., 2012; Zhao et al., 2015). Together, these studies indicate that electroactive microbes associated with EET are both phylogenetic and metabolic diverse, involving both Bacteria and Archaea.

Metabolic versality can provide unique insights into the unrecognized biogeochemical signals and the evolution of microbial metabolisms. Metabolic studies of microbial extracellular electron uptake has mostly been done on iron reducing bacteria affiliated with *Geobacter sp*.(Kumar et al., 2017; Ueki et al., 2018; Heidary et al., 2020) and *Shewanella sp*.(Rowe et al., 2018; Zou et al., 2019). Few studies extended to other functional guilds, such as iron oxidizers (Carbajosa et al., 2010; Summers et al., 2013; Doud and Angenent, 2014). It is hypothesized that iron oxidizers can perform EET due to their dense outer membrane and periplasmic anchored c-type cyctochromes in their native iron oxidation pathway (Croal et al., 2007; Jiao and Newman, 2007). Additional research is required to explore the bounds of EET metabolism in the tree of life and in various functional microbial guilds.

One of the widely applied BES type is known as aerobic biocathode, which accepts electrons from poised potential electrodes, while reducing O_2_ to generate energy (Ter Heijne et al., 2010). Aerobic biocathodes with mixed cultures were shown to increase the kinetic efficiency of graphite cathodes (Debuy et al., 2015), making biocathode performance similar to those with abiotic platinum based graphite cathodes (Cristiani et al., 2013). Furthermore, Rabaey et al. (2008) reported that mixed community based biocathodes are more efficient in terms of power than pure cultures due to the density and co-metabolic activity; *Sphingobacterium, Acinetobacter* and *Acidovorax* spp. were the dominant taxa of the aerobic biocathode community. Reimers et al. (2006) detected *Pseudomonas* as the dominant species at the aerobic biocathode of a microbial fuel cell on the ocean floor. *Alkalilimnicola, Bacteroidetes, Nitrosomonas* were detected as dominant taxa in aerobic biocathodes inoculated with municipal wastewater (Du et al., 2014). In a recent study, electroactive aerobic biocathode community enriched from aerobic activated sludge was dominated with *Nitrosomonas, Gordonia polyisoprenivorans, Nitratireductor* and *Spingobium*. Using metagenomics on biomass from an aerobic biocathode, originally inoculated with marine sediment, Wang and coworkers found that one of the dominant taxa in the mixed electroactive community was affiliated to *Chromatiaceae*a putative iron oxidizer (Wang et al., 2015). While these studies focused on the dominant taxa as the electroactive members, additional taxa reside in the cathode biofilms with relative abundances, ranging between 1% and 10%, making the electroactive member of a functional guild difficult to identify. As heterotrophic taxa utilizing secondary microbial products (Okabe et al., 2005) can co-exist on biocathodes together with electro-active autotrophic taxa, the need for a method to characterize species in terms of their energy source becomes more apparent.

We contend that further information regarding the taxa performing EET and the prevalence of EET metabolism is important, and functional guilds of ammonium (NH_4_) methane (CH_4_), thiosulfate (S_2_O_3_^2-^) and hydrogen (H_2_) oxidizers can perform EET, as they are known to use outer membrane cytochromes in their main oxidation pathway. The detection of EET metabolism within these guilds may also suggest an ability to switch between metabolisms and might help to unravel the evolution of EET metabolism on microbes. In addition, the detection of specific genera performing EET is central in our efforts to isolate pure electro-active cultures and elucidate EET mechanisms.

In this study, we first enriched H_2_, S_2_O_3_^2-^, or CH_4_ and NH_4_^+^ oxidizing guilds from a sand filter for groundwater treatment, where we observed a continuous inlet flow of ferrous iron, hydrogen sulfide, methane and ammonium (Gülay et al., 2016, 2019). We, then, examined the electrochemical activity and electroactive members of these enrichments with the aim to identify a microbial member of anticipated function performing extracellular electron transfer and to characterize their electron uptake efficiency. We also identified the electron energy distribution across investigated metabolisms and predicted taxa utilizing electron energy of the electrode and secondary metabolic products, using co-occurrence analysis of metagenomic libraries and abundance values.

## Material and Methods

### Sampling site

Sand-filter material for microbial members of anticipated functions performing extracellular electron transfer enrichments and 16S amplicon sequencing was collected from after-filters of Islevbro waterworks, as described in Gülay et al. (2016). The upper 0–10 cm of top filter layer was collected from three random positions in selected rapid sand filters (RSFs) treating groundwater for drinking water purpose. The rapid sand filter at Islevbro waterworks was chosen based on previous analysis of Gülay et al. (2016); where we observed a continuous groundwater inlet flow of ferrous iron, hydrogen sulfide, methane and ammonium (Gülay et al., 2016, 2019)

### Pre-enrichments

Sand-filter samples (approximately 3 g wet weight) were suspended in 5 ml minimal Wolfe’s mineral medium (MWMM) at pH 7 and cells were dislodged by shaking the mixture at 150 rpm for 30 min. The supernatant was used as inoculum for enrichment experiments. Briefly, 10 mL of aliquot was incubated at 60% maximum water-holding capacity in 120 ml serum bottles. Before inoculation, the pH of MWMM was adjusted to pH 7.5 with 1M NaHCO_3_. No additional organic carbon was added. The serum bottles were tightly capped with blue butyl stoppers and flushed with synthetic air (21% O_2_ in N_2_) for 5 min to ensure sterile oxic conditions. The serum bottles for methane and hydrogen oxidizers were amended with 60 mL of 100% methane and 100% hydrogen, respectively, by injection through the rubber stoppers. The microcosms of S_2_O_3_^2-^ and NH_4_^+^ were supplemented with 10 mM of NH_4_Cl, and S_2_O_3_^2-^respectively, as electron donors. Enrichments were incubated at room temperature in the dark with shaking. After an incubation period of 15 days, 10 mL of the enrichment cultures were used to inoculate new 120 ml serum bottles, containing 60 mL fresh aerobic MWMM media. The microorganisms were transferred to new media every ∼14 days for a period of 96 days before they were used in BES as inoculums.

### Bioelectrochemical reactor and operation

Two-chamber H-type BESs were constructed from two 250-mL plexiglass boxes joined with silicone rubbers. The two chambers were separated by a pretreated cation exchange membrane (CMI-7000S Cation Membrane, Membranes International INC.). Electrodes were carbon felt (3 cm length, 3 cm width, 1.12 cm thickness, 0.72 m^2^/g surface area, Alfa-Aesar, USA). Titanium wire (0.06 cm diameter; Sigma) at a length of 12 cm was polished with sand-paper to remove the oxide layer. The wire attached to the electrodes was then autoclaved. Reference electrodes (Ag/AgCl, Sentek, Geyer) was sterilized with 70% (v/v) ethanol in a laminar airflow cabinet. 220 mL of sterile basal MWMM medium was supplemented with 2 mL/L vitamin solution (ATCC ® MD-VS ^™^) and 2 mL/L trace mineral solution (ATCC® MDTMS^™^) before addition to single BES chambers. Both chambers were inoculated by addition of 20 mL pre-enriched culture (working volume = 220 mL; headspace = 30 mL). The headspace of all reactor chambers was flushed with sterile air to supply continuous mixing and O_2_. For chronoamperometric incubations, the BESs were connected to a potentiostat (Ivium Technologies, Ivium-n-Stat Multichannel potentiostat), that applied a cathode potential of −0.450 V (vs. AgCl). Each BES type (inoculated with enrichments of H_2_, S_2_O_3_^2-^, or CH_4_ and NH_4_^+^oxidizing microbes) was run in duplicate BES reactors. Parallel BES reactors were operated as controls without biomass. Identical voltage settings were used for both inoculated and controls reactors. No biomass growth was detected for both controls (1) that were abiotic and (2) that were not fed with any electron donor. Before inoculation of BES, all reactors produced negligible current, similar to or lower than the control experiments at the same potential.

After 10 days, cells attached to the cathode electrodes in the BES reactors were dislodged and collected as described in “Cell counts and DNA extraction” section. Collected cells were transferred into new BES reactors with media. At the end of BES operation (day 40), these BES reactors were subjected to cyclic voltammetry and then sacrificed for microbiological, molecular and chemical analysis.

### Cyclic voltammetry

At the end of the BES run (day 40), CV was applied to characterize the catalytic behavior of the biocathodes. The scan range of CV was from ⍰0.1 V to ⍰-0.5 V, with a scan rate of 1 mV/s, using the Ivium-n-Stat Multichannel potentiostat. We applied low, 1 mV/s, scan rate to minimize the background capacitive current and the kinetic limitations of interfacial electron transfer between the microbial cells and the electrode (Marsili et al., 2008). As selection of informative potential range is crucial in CV due to the harmful oxidizing or reducing conditions (Marsili et al., 2008); we applied −0.5V to 0.1V range, after testing that CV did not disturb the on-going current production.

### Chemical measurements

Liquid samples (5 mL) were taken at day 10 and 40 from the cathode compartment through a rubber septum, using a 10 mL syringe equipped with a sterile needle. Samples were immediately filtered through 0.22 **μ**m filters. The filtrate was used to measure dissolved organic carbon (DOC) concentrations. Samples without filtration were used for total organic carbon (TOC) analysis. TOC and DOC were determined by a total elemental carbon analyser (LECO CS□225).

### Cell counts and DNA extraction

At the end of the first BES run (10 days) and second BES run (40 days), the working electrodes were suspended into 15 mL cell-free 1X MWMM media and vortexed for 3 minutes to dislodge cells from the electrode. From each BES, cells in a 0.5 mL aliquot were stained with SYTO (SYTO9; Invitrogen, Carlsbad, CA, USA) and counted using a confocal laser scanning microscope equipped with an Ar laser (488 nm) ([CLSM] TCS SP5; Leica, Germany). Stained images were analyzed with a Leica AS AF Lite instrument (Leica, Germany) and ImagePro software (MediaCybernetics, USA).

DNA was extracted from 1) serum bottles with pre-enriched biomass, just before inoculation into BES, 2) after 11 and 36 days of incubation in the BES by using the FastDNA Spin kit (MP Biomedicals, Solon, OH, USA) and the manufacturer’s instructions. The concentration and purity of extracted DNA were checked by a NanoDrop 2000 spectrophotometer (NanoDrop Technologies, Wilmington, DE, USA).

### Efficiency calculations

Current density was calculated according to the equation:

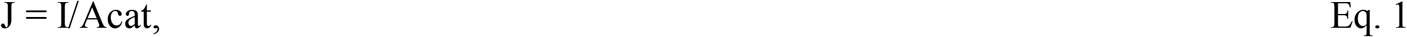

where J is the current density in A/m^2^, I is the current (A) and A_Cat_ is the cathode surface area (m^2^).

The energy consumed was calculated by the following equation:

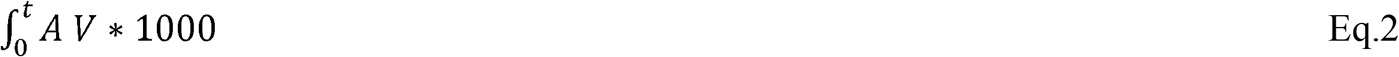

with I the current (A), V the voltage (V), and t time (s). The abiotic energy loss in the BES set-up was calculated with Eq.2 using the measurement of the control experiment.

The energy captured in cellular biomass BES runs were estimated by the converting the cell counts (using the Stoichiometry.1) to Gibbs free energy using the following stoichiometry (Liu et al., 2016):

#### Biomass formation

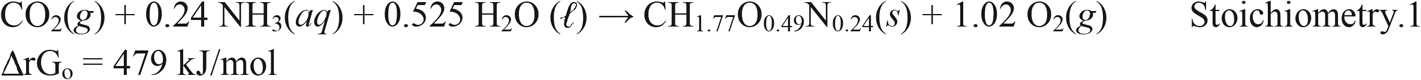

The ΔrG_o_ value for biomass is based on the report that the Gibbs free energy of formation of biomass in *Escherichia coli* is –46 kJ/mol carbon (Liu et al., 2016).

The energy used to produce particulate and dissolved organics in BES was estimated from the TOC-DOC and DOC values, respectively. The energy consumed for particulate carbon was estimated using Stoichiometry.2 assuming all products are de-attached cells from the biofilm. Carbohydrate (CH_2_O) was chosen as the dissolved carbon product assuming measured DOC values are CH_2_O. The following stoichiometry was used to estimate the energy cost:

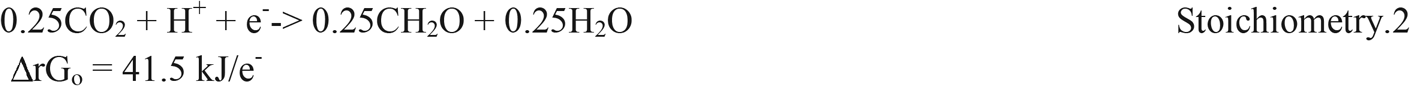

### 16S Amplicon sequencing and bioinformatic analysis

PCR amplification and 16S amplicon sequencing were performed at the DTU Multi Assay Core Center (Kgs Lyngby, DK). Briefly, extracted DNA was PCR amplified using 16S rRNA bacterial gene primers PRK341F (50-CCTAYGGGRBGCASCAG-30) and PRK806R (50-GGACTACNNGGGTATCTAAT-30), targeting the V3 and V4 region (Yu et al., 2005). PCR products were purified using AMPure XP beads (Beckman-Coulter) prior to index PCR (Nextera XT, Illumina) and sequenced by Illumina MiSeq.

Raw 16S rDNA amplicons from MiSeq were denoised with Deblur, processed and classified with QIIME and Mothur (Schloss et al., 2009; Caporaso et al., 2010). Chimera checking was performed with the software UChime (Edgar et al., 2011). High quality sequences were clustered at 100% evolutionary similarity.

The differential absolute abundance value for each genus was calculated by transforming the relative abundance values from 16S rRNA libraries to absolute abundance values, using the total cell counts obtained from at the start (day 0) and at the end (day 40) of BES operation.

### Network analysis

For network inference, the OTU libraries of BES at day 40 were used to calculate co-occurrence maps. Taxa with a relative abundance below 1% were trimmed from the OTU library. All possible Pearson linear correlations between OTUs were then calculated. For co-occurrence analysis, a valid co-occurrence event was considered to be a robust correlation if the Pearson correlation coefficient (r) was both larger than 0.6 and statistically significant (P-value 0.05) (Steinhauser et al., 2007). Only positive correlations were selected for further analysis. Top 5 genera with the highest absolute abundance difference (Figure 3A) between day 0 and day 40 were tentatively labeled as primary consumers of electrons from the cathode. Genera that co-occurred with these primary consumers in each of the BES runs were selected and labelled as secondary consumers. For each BES type, co-occurred taxa with secondary consumers were selected and labelled as “tertiary consumers”. The nodes of the network identify the genera, with the node size showing their average relative abundance of those genera, and the edges were used to show a strong, positive, and significant correlation between nodes.

## Results and Discussion

### Autotrophic enrichments

Inocula from a rapid sand-filter were used to enrich microbes on H_2_, S_2_O_3_^2-^, or CH_4_ and NH_4_^+^, as the sole electron donor, respectively. All cultures displayed visible biomass after 3^rd^ transfer (42 days of incubation). *Methylomonaceae* and *Methylophilaceae* dominated the enrichments amended with CH_4_, *Xanthobacteriacea* and *Weeksellaceae* dominated the H_2_ fed enrichments, *Hydrogenophilaceae* and *Streptococcaceae* dominated NH_4_^+^ fed enrichments (Figure 3B). Enrichments amended with CH_4_ were dominated by well-known methane oxidizers; these taxa were previously detected from CH_4_ fed column reactors inoculated with sand-filter material (Papadopoulou et al., 2019). Dominant taxa in all enrichments yield over 50% of total community abundance, indicating that they outcompeted the other members, performing the anticipated function. However, no known nitrifiers were observed in NH_4_^+^ fed enrichments.

### BES operation and biocathodic testing of enrichment cultures

After inoculation of the BES with the different enrichments, the current density of duplicate BES increased slowly (data not shown). Transfer of the biomass after 10 days of operation to a new cathode chamber leads to a significant increase in current density (Supplementary Figure 1). Immediately after transfer, S_2_O_3_^2+^enrichments increased its current density in both BES duplicates. The enriched NH_4_^+^ and CH_4_ oxidizers reacted slower. First after 16 days, one of the duplicate BES with enriched CH_4_ and NH_4_^+^ oxidizers increased their current density (Figure 1). The current density reached around −0.03 (rep 1) and −0.06 (rep 2), −0.04 (rep 1) and −0.07 (rep 2), −0.09 (rep 1) and −0.07 (rep2), −0.07 (rep1) and −0.05 (rep 2) I/A.m^-2^ for H_2_, S_2_O_3_^2-^, or CH_4_ and NH_4_^+^ enrichments, respectively.

**FIGURE 1.**
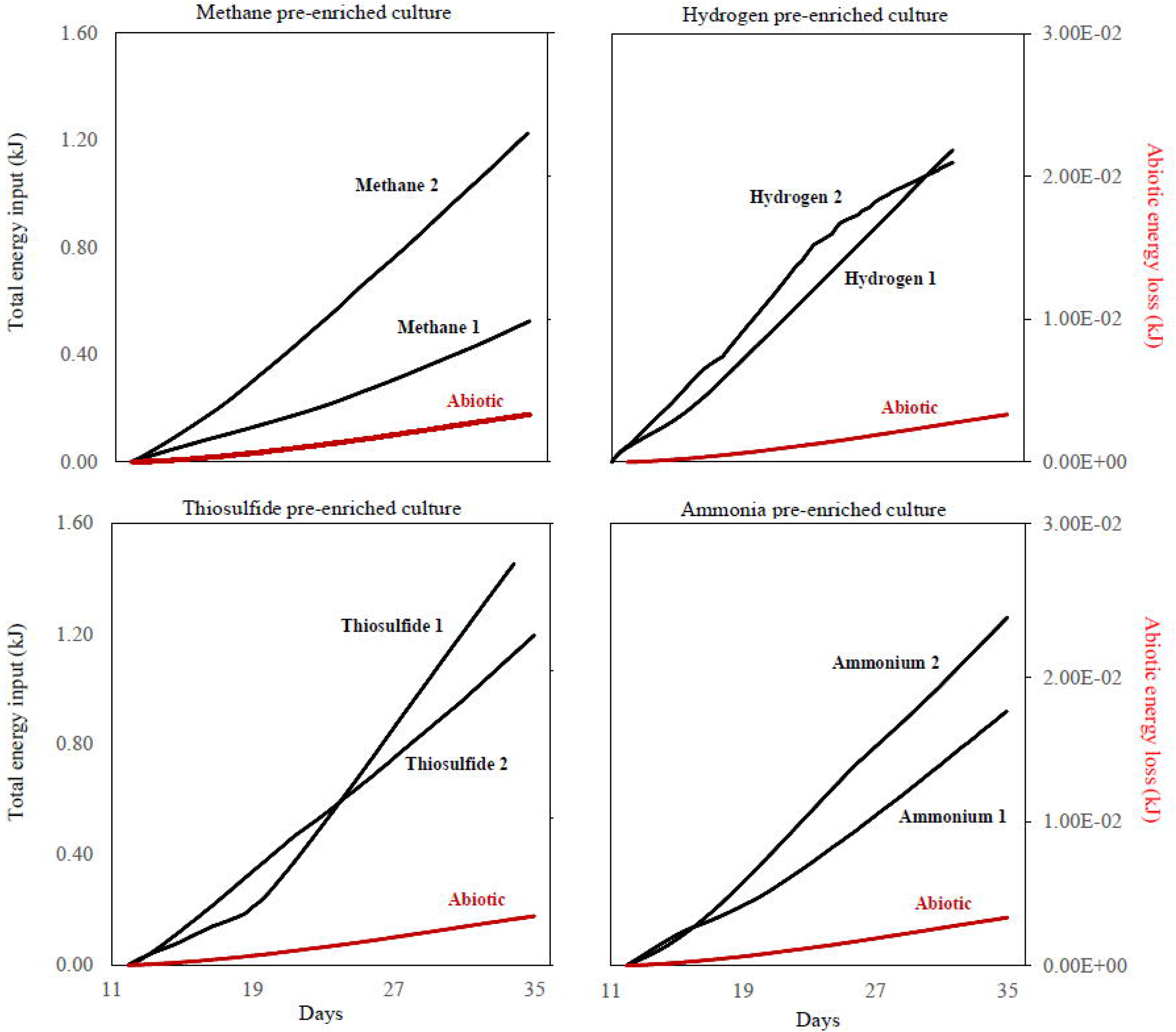
Total energy (kJ) given (black) in biocathodes with methane, hydrogen, thiosulfide and ammonia pre-enrichments and (red) energy loss calculated from the control reactor without inoculation.

However, none of the BES with CH_4_ and NH_4_^+^ BES reached their maximum current density, as the current were still increasing at the time of BES closure (Supplementary Figure 1). Although H_2_ and S_2_O_3_^2-^ enrichments showed higher maximum current density than CH_4_ and NH_4_^+^enrichments, all BES incubations developed a current with time indicating biological growth and activity (Figure 1). A visible biofilm was formed at the surface of the electrodes in all BES types at day 40 (Supplementary Figure 2). BES with H_2_ enrichment reached a maximum current density of −0.09 A/m^2^ and −0.07 A/m^2^ on day 24 and 29 for replicate 1 and replicate 2, respectively. The BES with thiosulfate enrichment (replicate 2) reached a maximum current density of −0.097 A/m^2^ on day 26, while replicate 1 did not reach its maximum current density.

The dislodgement of electrode-attached communities and allowing their re-attachment to a new electrode after 10 days of BES operation might have enabled a faster start-up of BES reactors due to a possible pre-selection of taxa capable of accepting electrons from the cathodes. BES needed several weeks before their maximum current was reached, in line with other microbial oxygen biocathode studies, suggesting that dominant taxa in biocathode community consist of slow-growing autotrophic organisms.

### Catalytic Behavior for Oxygen Reduction

Cyclic Voltammetry (CV) is a method to detect redox active molecules by sensing the potential difference across the interface of electrode and biofilm and thus allows one to identify electrochemical reactions occurring at the electrode surface (Rusling et al., 2008).

The maximum current density of −245 −1000 mA was reached at a cathode potential of −0.5V vs Ag/AgCl for replicate BES with methane enrichments. Similarly, at a cathode potential of −0.5V vs Ag/AgCl, −650 mA and −390mA, −395mA and −520 mA, and −320 mA and −585 mA were reached for replicate BES with H_2_, S_2_O_3_^2-^ and NH_4_^+^enrichments, respectively. At more positive potentials, the current was stable, except for BES with hydrogen enrichments. A cathodic peak ranging from −60 mV (replicate 2) to −100 mV (replicate 1) was detected at a cathode potential of 0 V vs Ag/AgCl. When comparing the CV results of operated BES together with the CV obtained for chemical oxygen reduction (red profile; Figure 2), indeed the oxygen reduction reaction was catalyzed by the microbes. The presence of biofilm resulted in a current multiple time higher than the current from chemical oxygen reduction (Supplementary Figure 3). At negative currents (below-0.32V vs Ag/AgCl), maximum current achieved by all BES types exceeds the maximum current reached with abiotic control.

**Figure 2.**
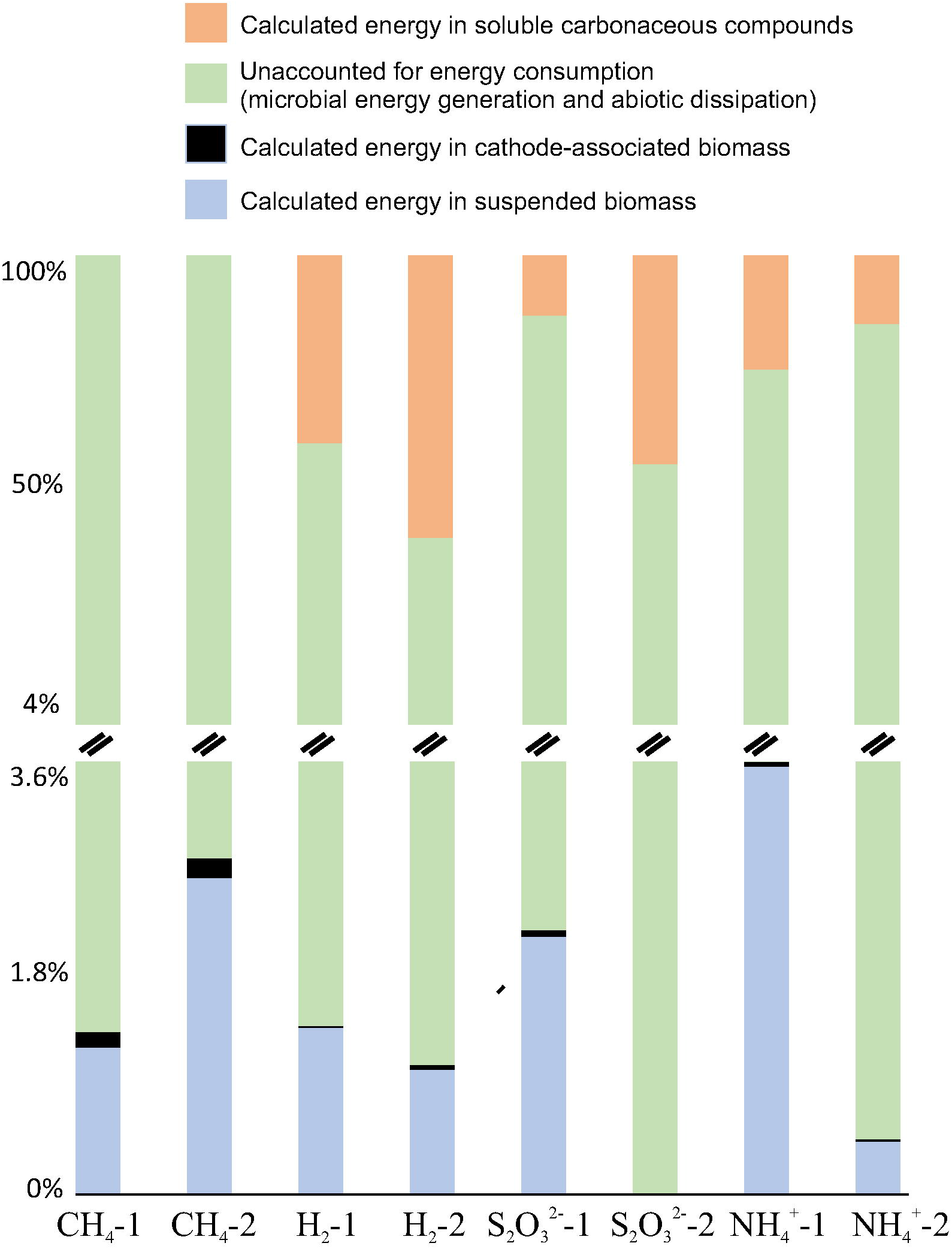
Distribution of energy (%) consumed by biofilm on electrodes of BES inoculated with methane, hydrogen, thiosulfide and ammonia pre-enrichments. The green bars on the upper corner represents energy directed to cathode-associated biomass.

The peak potentials in this study were similar to those observed by Xu et al., 2010, where cathodic biofilms were enriched by immersing electrodes in natural seawater. Their CV results yielded reduction peaks at around −360 to −460 mV vs. Ag/AgCl. In reported CV profiles, the potential that *Pseudomonas aeruginosa* reached its maximum current (−400 mV vs. Ag/AgCl) at the cathodic part (Cournet et al., 2010) and the electrochemical behavior of multi-heme cytochromes (cytochrome c3) of the sulfate-reducing bacteria *Desulfomicrobium norvegium* (Sallez et al., 2000) were in line with our results. On the other hand, the electrochemical behavior of outer membrane cytochromes OmcA and MtrC purified from *Shewanella oneidensis* MR-1 − reaching their maximum current peak at −300 V Ag/AgCl − were not observed in CV profiles (Meitl et al., 2009), suggesting that known EET (mtrCAB) pathway is not the dominant EET pathway in our study.

### Energy consumption and energy distribution of BES communities

The total energy consumption in all BES reactors and abiotic controls were calculated at the end of the experimental period to compare the energy uptake efficiency of the different enrichments (Figure 1). We further estimated the distribution of the dissipated energy into different fractions, including biofilm associated cellular material, suspended cellular material, and dissolved carbonaceous compounds (Figure 2).

In contrast to the similarities in the energy uptake profiles (Figure 3A), the energy distribution was distinct across the BES community. The largest energy directed to biomass growth was detected for the methane enrichment, followed by a replicate of S_2_O_3_^2-^ enrichment. The largest energy directed to soluble microbial products (SMP) was observed for hydrogen enrichment, and apparently no energy was directed to SMPs for methane enrichments. Furthermore, the BES reactors with ammonia and methane enrichments showed the largest energy directed to suspended material (including de-attached cells and the attached extracellular polymeric substances).

**Figure 3.**
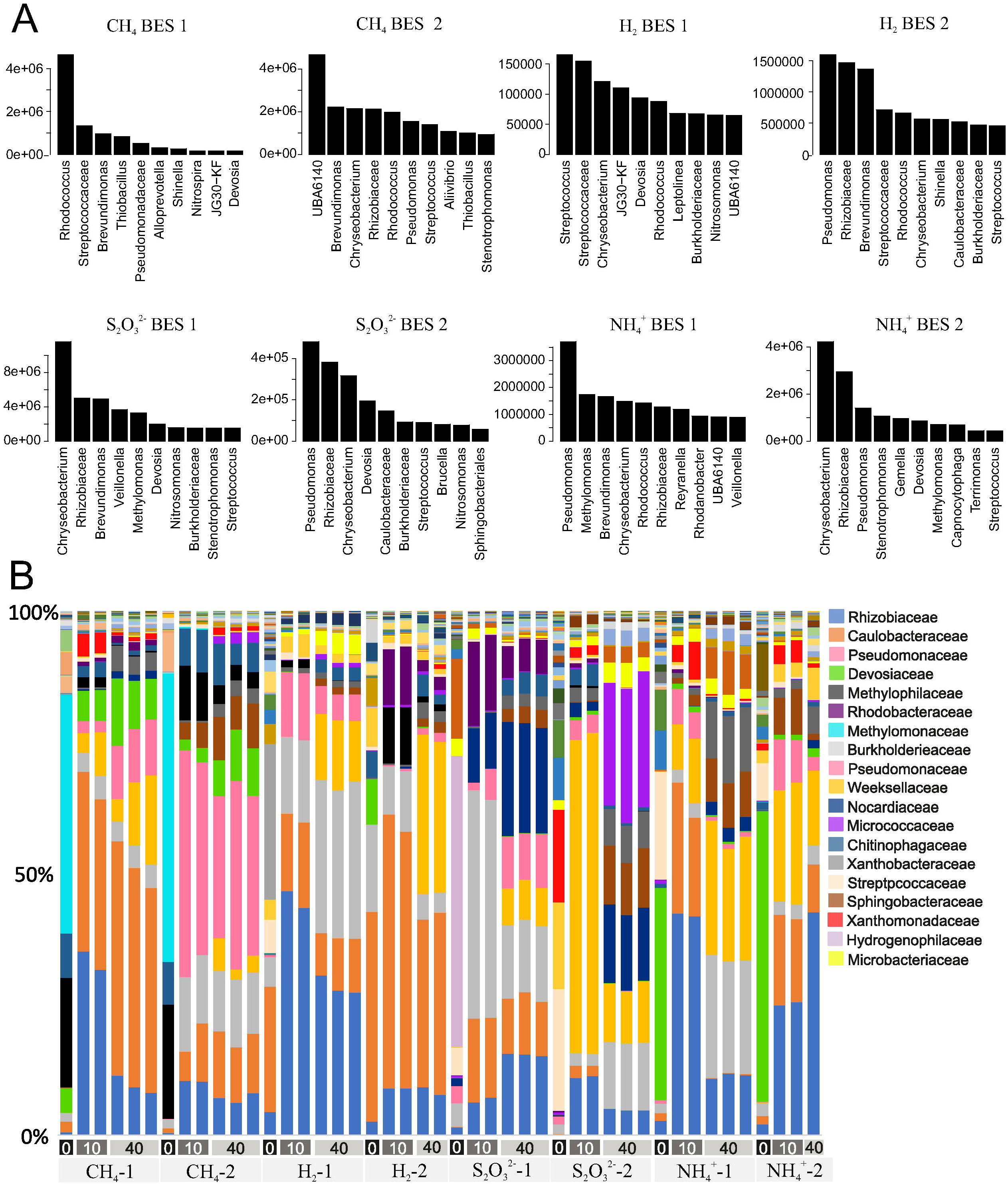
(A) Changes in absolute abundance of genera in the cathode-associated community BES between day 0 and day 40. The top 10 genera (averaged over the replicates of a stage) having the highest level of abundance change are shown. (B) Relative abundance and family level taxonomic classification of 16S rRNA amplicons across all BES. For each BES type, microbial composition of day 0, day 10, and day 40 are plotted along the x axis.

Rittmann and colleagues (2001) reported that sulfide oxidizing bacteria directed an energy of 0.008 kJ/gH_2_S to cell growth, while performing sulfide oxidation coupled to oxygen reduction. This was an order of magnitude higher than the energy that sulfide oxidizers directed to cell growth while performing EET in our study. Likewise, aerobic nitrifiers was reported to allocate 0.0004 kJ/gNH_4_^+^ of their energy to cell growth during ammonia oxidation, which is higher than the energy directed while activating their EET pathway.

### Microbial community structure

The cathode graphite electrodes showed a dense packed biofilm (Supplementary figure 2). The 16S rRNA amplicon sequencing showed a significant shift in microbial taxa from the day of BES inoculation (day 0 represented by the pre-enrichments in serum bottles) to the end of BES operation (day 40). (Figure 3B).

The relative abundance ratios were combined with the cell density counts of samples taken at the start and the end of the BES operation. This enabled us to estimate the cell densities of each genus on day 0 and 40 to identify taxa growing at the electrode (Figure 3A). Detected genera in replicate BES reactors were different in all BES types, as expected (Figure 3A). *Chryseobacterium, Brevundimonas, Pseudomonas* and unclassified members of *Rhizobiaceae* were detected in duplicate BES inoculated with S_2_O_3_^2-^ enrichments (Figure 3A). *Brevundimonas* were present in all BES types at cell densities ranging from 1.5 × 10^5^ to 5 × 10^6^ cells/reactor. In BES inoculated with methane enrichments, *Rhodococcus* and *UBA6140* showed 5-fold higher cell growth. The vast growth of UBA6140, genus of the *Methylophilaceae* - a known methane oxidizing clade - was in line with our hypothesis that methane oxidizers can perform EET, yet the ability of EET within *Methylophilaceae* has not to the best of our knowledge been reported before.

In BES inoculated with hydrogen enrichments, *Streptococcus, Pseudomonas*, unclassified members of *Streptococcaceae* and *Rhizobiaceae* were identified as the genera with highest growth. Although *Pseudomonas, Chryseobacterium* and *Brevundimonas* were detected at cell densities ranging from 1.5 × 10^5^ to 4 × 10^6^ - similar to the other BES types - *Methylomonas* and *Stenotrophomonas* were identified as taxa with active growth only in BES with NH_4_^+^enrichments.

Surprisingly, our results suggest that *Methylomonas, UBA6140*, and *Nitrosomonas* can uptake electrons from the cathode. While methane oxidizing taxa, *Methylomonas* and *UBA6140* has not been reported as electro-active taxa before, *Nitrosomonas* was also detected as a dominant taxon in other biocathode studies (Du et al., 2014; Mani et al., 2020). Further studies are required to confirm and examine mechanism of EET of CH_4_ and NH_3_ oxidizers.

Based on relative abundances in Figure 3B, Microbial characterization at day 0 revealed that the known methane oxidizers within the families of *Methlomonaceae* and *Methylophilaceae* were the most dominant methane oxidizers in methane enrichments. Further, *Xanthobacteriacea*, and *Caulobacteriaceae* was dominating in hydrogen enrichments, while *Hydrogenaphilaceae, Streptococcaceae Xanthomonodaceae* was abundant in thiosulfate enrichments. In ammonium enrichments, *Devosiaceae* were detected as dominant taxa with the relative abundance of 25% to 70%. *Rhizoniaceae* and *Caulobacteraceae* were became dominant in almost all BES on day 40, suggesting that they have high affinity for EET, and that their growth was not restricted by the initial microbial community composition.

Although *Methylomonaceae* were the dominant methane oxidizers in methane enrichments, they disappeared from both BES with methane enrichments at the day 40. On the other hand, *Methylophilaceae* remained dominant. While the members of *Devosiaeae* were dominant in ammonium and methane fed pre-enrichments, they increased dominance in the BES with methane oxidizing enrichments. *Nocardiaceae* and *Rhodobacteriaceae* increased in abundance in BES with thiosulfate enrichments after 11 and 40 days. Also, *Weeksellaceae* increased in abundance after 40days of BES operation with thiosulfate enrichments. The microbial community in BES with hydrogen and thiosulfate enrichments were dominated by of *Microbacteriaceae*, while *Xanthobacteriaceae* and *Rhodobacteriaceae* were abundant in BES inoculated with ammonium enrichments.

### Trophic levels of BES communities

There are no obvious genetic markers to detect EET pathways (Wang et al., 2015). Therefore, we explored the communities in the BES initiated with H_2_, S_2_O_3_^2-^, CH_4_ and NH_4_^+^enrichments by amplicon sequencing in an effort to link EET physiology to taxonomy.

The top 5 taxa with highest increase in cell numbers between day 0 and 40 were considered to have electroactive and autotrophic lifestyle. However, taxa having lower abundances than these top 5 taxa were expected to perform direct interspecies electron transfer or oxidation of organic products that were generated by autotrophic taxa, making it difficult to predict their source of energy. Here, a framework using co-occurrence analysis and microbial abundance patterns was applied to identify the source of energy for taxa having low abundance. Co-occurrence networks been used to describe the complex pattern of interrelationships between OTUs (Barberán et al., 2012; Faust and Raes, 2012) and its topological properties are very informative to identify taxa having mutualistic and commensalistic relationships (Newman, 2003; Meur and Gentleman, 2012; Schich et al., 2014).

To assess non-random co-occurrence patterns, we first combined community data at day 40 from all BES and selected the co-occurring taxa, using network inference based on significant and strong positive correlations (using non-parametric Pearson; Pearson and Lipman, 1988). Restricting the analysis to only those taxa showing significant relationships, the correlation score, on average, increased to 0.78 and P-Value of 0.002. Once we determined that the co-occurrence patterns were non-random, we merged the co-occurrence network with absolute abundance scores of each genus in each BES. Co-occurred taxa with those having the highest cell growth were considered to secondary consumers, scavenging the energy left by autotrophs that consumes electron energy released from the electrode.

The BES network was comprised of highly connected OTUs structured among densely connected groups of nodes of dominant taxa and forming a clustered topology (Figure 4). Unclassified members of *Streptococcaceae* and *Rhizobiaceae, Nitrosomonas, Methylomonas, Streptococcus, Brevundimonas, Chryseobacterium, Pseudomonas, Rhodococcus* and *UBA6140* were predicted as taxa performing extracellular electron uptake. Surprisingly, members of *Candidatus Nitrotoga, Nitrospira* and *Nitrobacter*, which are all known nitrifiers having obligate chemoautotrophic lifestyle, tended to co-occur more than expected by chance with *Brevundimonas* and *Pseudomonas* that were identified as electroactive autotrophs in this study. In our study, they were detected as secondary consumers suggesting a metabolism different than heterotrophy. It is possible that these taxa may do direct intercellular electron uptake from the autotrophs. 21 more genera were identified as taxa that may utilize secondary metabolites or electrons from the taxa utilizing direct electron energy at the cathode biofilm (Table 1).

**Table 1:** Identification of genera having different source of energy predicted by co-occurrence analysis and abundance patterns.

| Level 1 |  | Level 2 |  |  |  |
| --- | --- | --- | --- | --- | --- |
| Source of energy:<br>electron energy |  | Source of energy:<br>secondary electron energy or energy from microbial products |  |  |  |
| Brecundimonas | Pseudomonas | Aminobacter | Nitrospira | Phreatobacter | Stenotrophomonas |
| Caulobacteraceae | Rhizobiaceae | Candidatus_Nitrotoga | Nocardioides | Pigmentiphaga | Weeksellaceae |
| Chrysebacterium | Rhodococcus | Chitinophagaceae | Nordella | Ralstonia | Xanthomonadaceae |
| Devosia | Streptococcus | Curvibacter | OM190 | Reyranella |  |
| Gemelia | Thiobacillus | Dietzia | Paracoccus | Sphingobacterium |  |
| JG30-KF | UBA6140 | Ferruginibacter | Pedosphaeraceae | Sphingopyxis |  |
| Methylomonas | Vellionella | Nitrobacter | Phenylobacterium | Sphingorhabdus |  |

**Figure 4.**
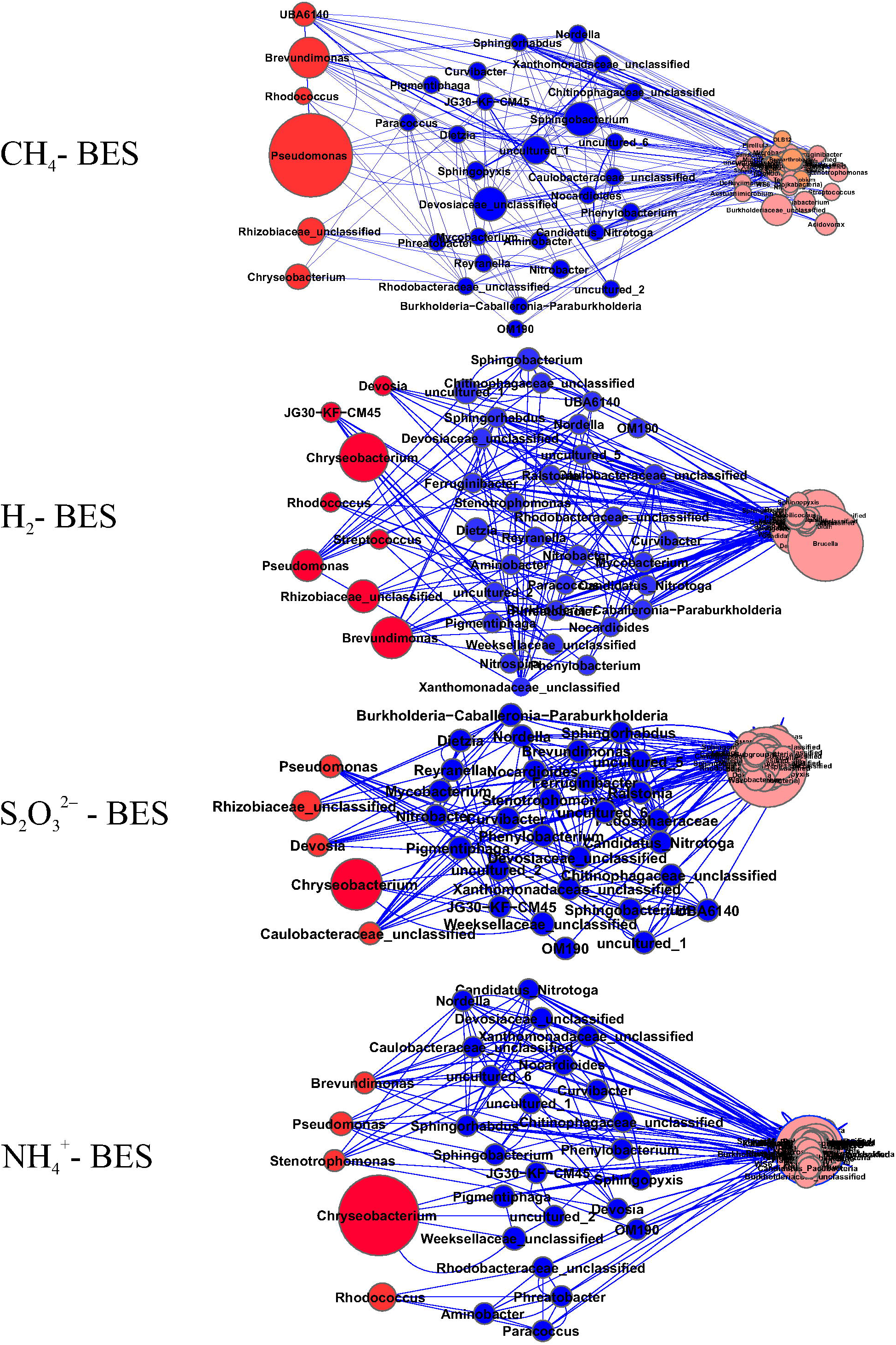
Network of co-occurring genera of all BES based on correlation analysis. A connection stands for a strong (Pearson’s R>0.6) and significant (P-value <0.05) correlation. The size of each node is proportional to its relative abundance.

## Acknowledgements

This research was financially supported by NovoNordisk Foundation (Project: BIOCAT-Novel Bio-cathodic Microbes and their electron transfer pathways for Efficient Energy Harvesting).

