## Supplementary Material for "Different autotrophic enrichments yield efficient inocula for biocathodic applications"

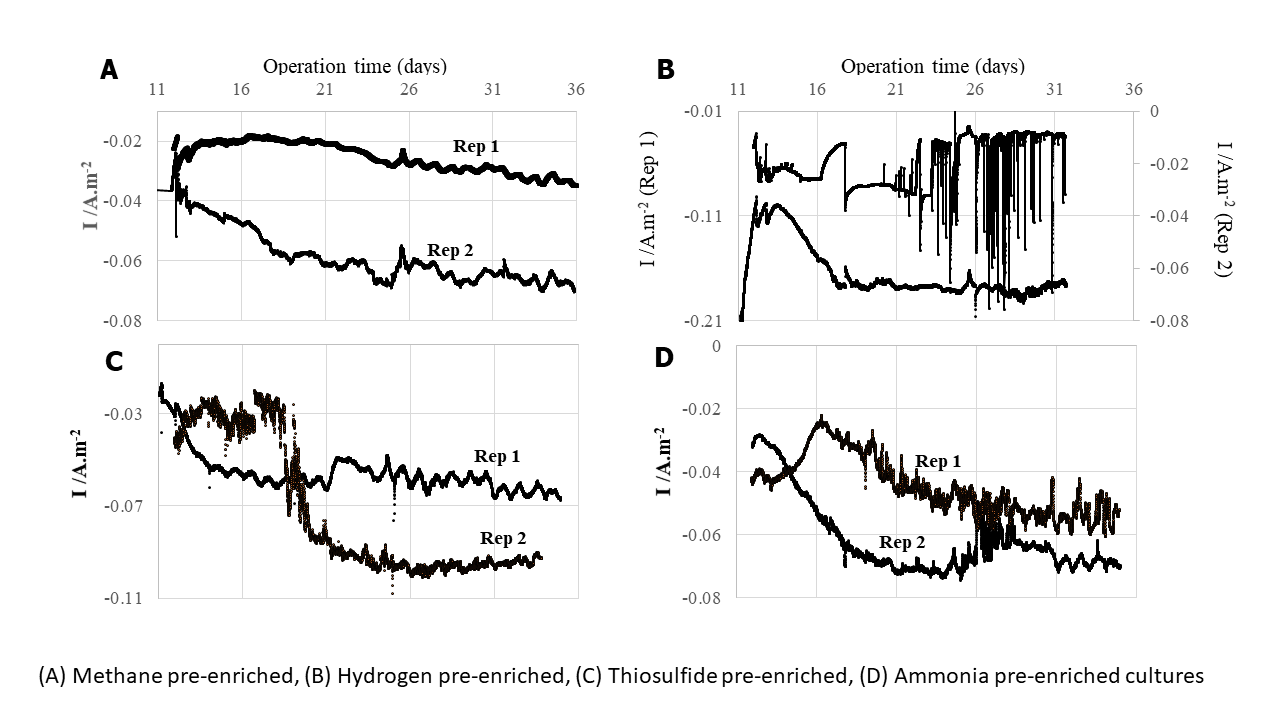

**Supplementary Figure1** Current density of biocathodes inoculated with methane, hydrogen, thiosulfide and ammonia enrichments. The chronoamperometric profiles were given starting from the second run (day 10 to day 40).

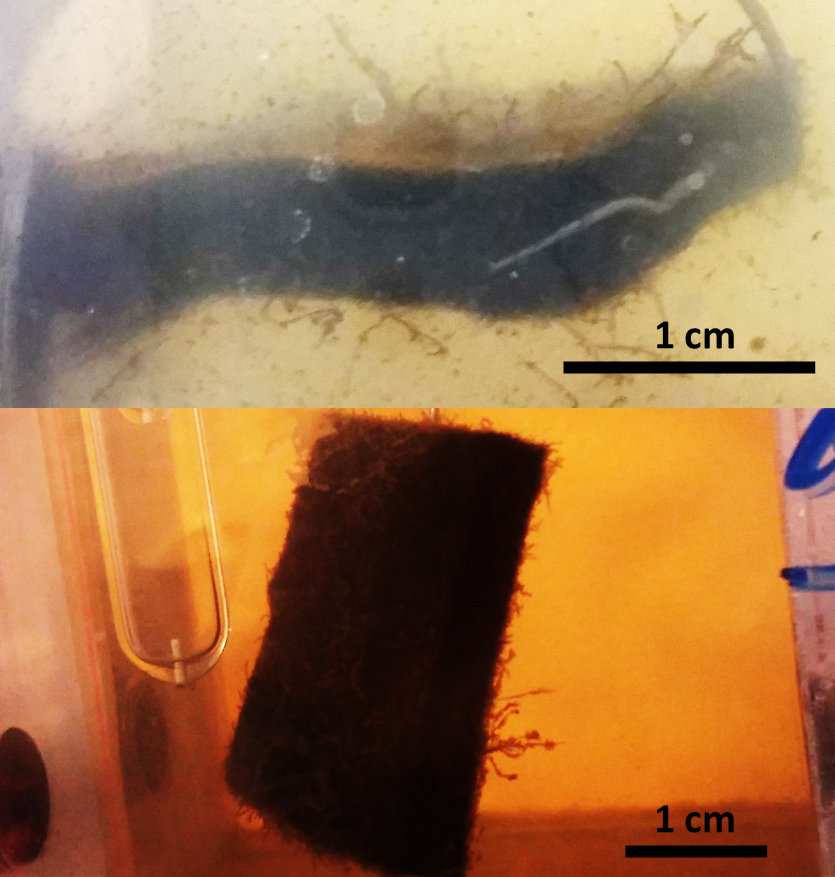

**Supplementary Figure 2** Biofilm developed at the surface of the electrode from BES inoculated with enrichments of ammonium and hydrogen. The picture was taken on BES operation day of 40 before the sampling.

**1 cm**

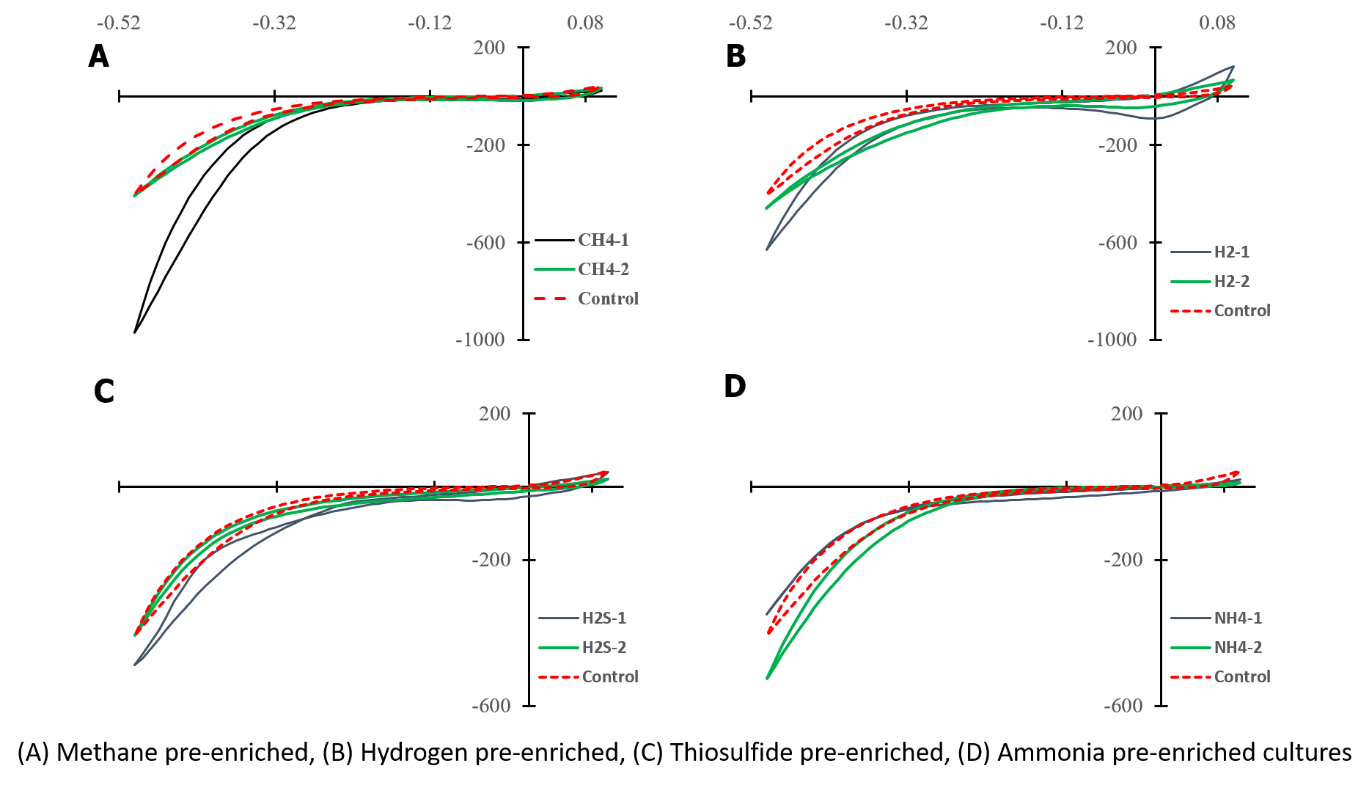

**Supplementary Figure3** Cyclic voltammetry (CV) graphs of biocathodes inoculated with methane, hydrogen, thiosulfide and ammonia enriched populations. CV was performed at the end of BES operation (day 40).
